# The dynamic transition of persistence towards the VBNC state during stationary phase is driven by protein aggregation

**DOI:** 10.1101/2021.02.15.431274

**Authors:** Liselot Dewachter, Celien Bollen, Dorien Wilmaerts, Elen Louwagie, Pauline Herpels, Paul Matthay, Ladan Khodaparast, Laleh Khodaparast, Frederic Rousseau, Joost Schymkowitz, Jan Michiels

## Abstract

Decades of research into bacterial persistence has been unable to fully characterize this antibiotic-tolerant phenotype, thereby hampering the development of therapies effective against chronic infections. Although some active persister mechanisms have been identified, the prevailing view is that cells become persistent because they enter a dormant state. We therefore characterized starvation-induced dormancy in *Escherichia coli*. Our findings indicate that dormancy develops gradually; persistence strongly increases during stationary phase and decreases again as persisters enter the viable but nonculturable (VBNC) state. Importantly, we show that dormancy development is tightly associated with progressive protein aggregation, which occurs concomitantly with ATP depletion during starvation. Persisters contain protein aggregates in an early developmental stage while VBNC cells carry more mature aggregates. Finally, we show that at least one persister protein, ObgE, works by triggering aggregation and thereby changing the dynamics of persistence and dormancy development. These findings provide evidence for a genetically-controlled, gradual development of persisters and VBNC cells through protein aggregation.

## Introduction

Bacteria can overcome antibiotic insults in different ways. Besides the well-known resistance phenomenon, bacteria can also survive antibiotic treatment by entering a transient antibiotic-tolerant state, called persistence. Persister cells are defined as rare phenotypic variants that survive antibiotic treatment, although they cannot grow in the presence of the antibiotic (1). However, since they can resume growth when the antibiotic pressure drops, persisters are assumed to contribute to treatment failure and the chronic nature of many bacterial infections (2, 3).

Although some active persister mechanisms have been identified (4–6), the prevailing view is that cells become persistent because they enter a dormant state (2, 3, 7). Indeed, many reports have shown that persisters are highly enriched in fractions of the population that are non-growing or slowly growing (8–10), have low ATP concentrations (11, 12) and/or low levels of protein expression (13). Recently, it was suggested that this dormant persister state is caused by the aggregation and concomitant inactivation of many important cellular proteins (14). Previous findings indeed suggest that persistence and protein aggregation are linked. Persister cells more often contain protein aggregates and artificially inducing or preventing protein aggregation affects persistence accordingly (14–16).

Besides persistence, other states of bacterial dormancy have been described. Among these is the so-called viable but nonculturable (VBNC) state; a phenotype in which cells have lost the ability to grow on conventional media that otherwise support their proliferation, even though they remain viable. Therefore, these VBNC cells cannot readily resume growth when provided with fresh nutrients but instead require specific conditions to resuscitate (17–19). Important similarities between persisters and VBNC cells exist; both are dormant bacterial phenotypes and both are tolerant to high antibiotic concentrations (1, 17, 18, 20). Because of these similarities, the hypothesis that persisters and VBNC cells are conceptually similar is gaining momentum (14, 17, 18, 21). According to this hypothesis, persisters and VBNC cells represent different developmental stages of the same dormancy program, where VBNC cells have obtained a deeper dormancy. However, detailed experimental evidence of the dynamic nature of persister formation and VBNC transition as well as genetic and molecular mechanisms to support this hypothesis are currently lacking.

We here confirm that a tight link between protein aggregation and persistence exists. Moreover, by examining the dynamics of protein aggregation in relation to the development of both persistence and the VBNC state, we provide compelling support for the hypothesis that persister cells can obtain a deeper dormancy and transition into the VBNC state. Our findings indicate that, during starvation, this transition is most likely driven by progressive protein aggregation which occurs concomitantly with ATP depletion. Persisters contain protein aggregates in an early developmental stage while VBNC cells carry more mature aggregates. Additionally, we show that persistence is a highly dynamic phenomenon and that persister levels vary strongly throughout stationary phase. Our data therefore indicate that the single-time-point investigation of persistence that is current standard practice is insufficient to fully capture this dynamic phenotype. Finally, we demonstrate that at least one known persister protein works by accelerating protein aggregation and thereby changing the dynamics of persister development.

### Results

#### Cells enter the viable but nonculturable state in late stationary phase

To gain more insight into bacterial dormancy, we studied the behavior of *E. coli* in stationary phase; the growth phase where dormancy sets in (22). We incubated bacterial cultures for a period of 72 hours and monitored the number of colony forming units (CFUs) as well as the amount of viable cells in the population (Figure 1A). Viable cells were quantified by flow cytometry after staining with the membrane-impermeable viability dye SYTOX Green and enumerating the SYTOX Green negative cells (representative flow cytometry graphs shown in Figure S1A) (17, 19, 20). Viability at the latest time point was confirmed using RedoxSensor Green (RSG), an indicator for metabolic activity (Figure S1B) (8, 23). As expected, CFUs start to decrease in long-term stationary phase. However, the decrease in the number of viable cells is much smaller, indicating that cells adopt the viable but non-culturable (VBNC) state. Indeed, several bacterial species, including *E. coli*, are known to transition into the VBNC state upon nutrient starvation (17, 19).

**Figure 1:**
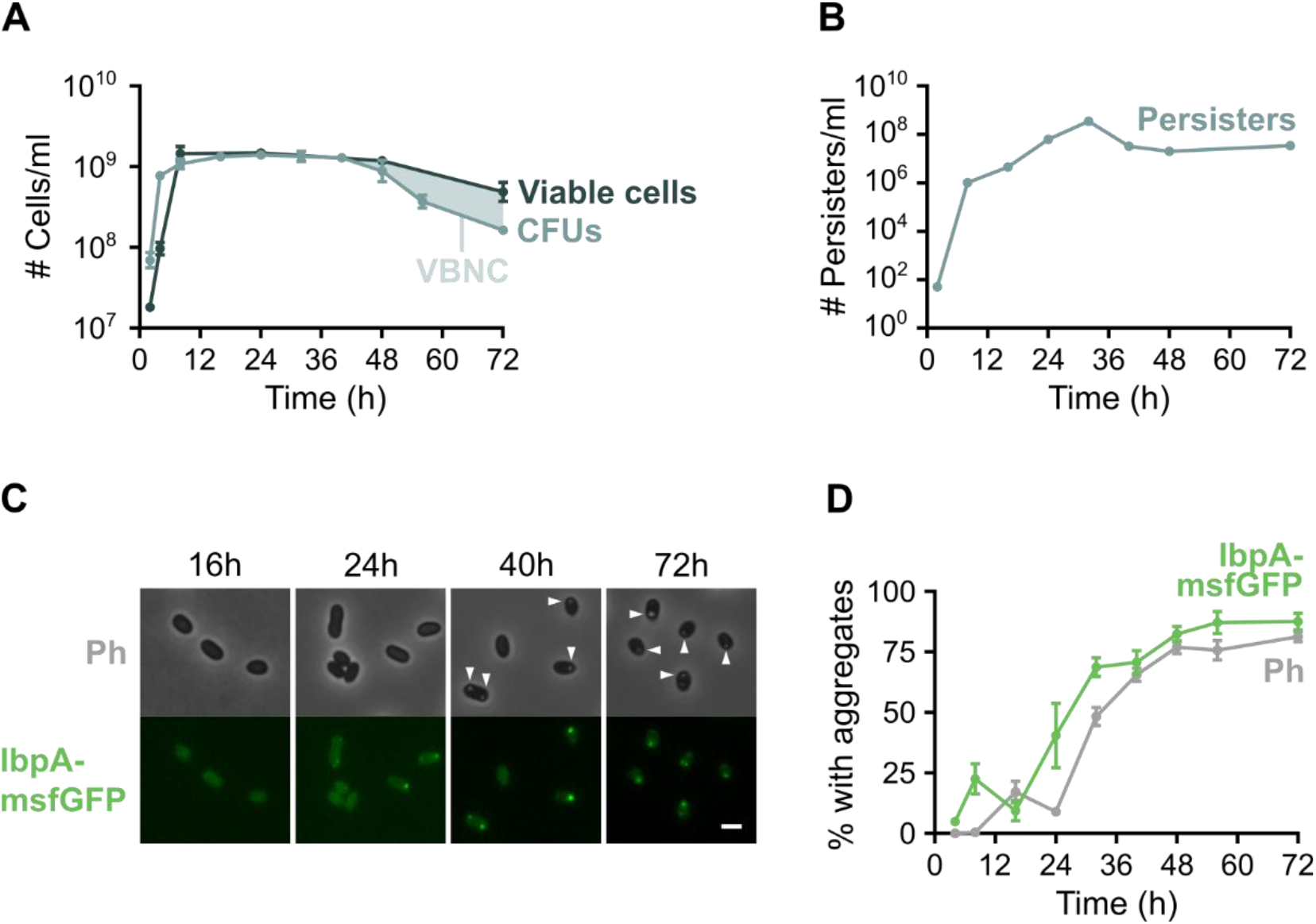
The VBNC state, persistence and protein aggregation are natural phenomena that occur in stationary phase. A) At different time points, the number of CFUs/ml of an *E. coli* culture was measured as well as the number of viable cells by counting SYTOX Green-negative cells with flow cytometry. The difference between both curves is the number of VBNC cells present in the culture. B) The absolute number of persister cells after treatment with ofloxacin was followed over time (error bars are too small to be visible). C) Microscopy images (phase contrast and GFP channels) of *E. coli ibpA-msfGFP* at different time points show the development of protein aggregates over time. Arrowheads point to aggregates that are visible in the phase contrast channel. Scale bar, 2 μm. D) Quantitative analysis of microscopy images was performed to determine the percentage of cells that carry protein aggregates. Aggregation was evaluated by the presence of fluorescent IbpA-msfGFP foci and phase-bright structures. For every repeat and every time point at least 50 cells were analyzed. Data are represented as averages ± SEM, n ≥ 3. Ph, phase bright.

#### Persistence is a highly dynamic phenomenon that is induced in stationary phase

To investigate the dynamics of persister development, we monitored the absolute number of persister cells over time. Cultures were treated with ofloxacin at several different time points before and during stationary phase and the number of surviving persister cells was measured (Figure 1B). As expected, the number of persister cells increases drastically upon entry into stationary phase. However, persistence is also highly dynamic during stationary phase, reaching a maximum after 32 hours of incubation. At this time point, approximately 25 % of all cells have become persistent. This percentage is much higher than what is usually recorded (typically between 0.001% and 1% of the population (2)) and demonstrates that persister fractions can be strongly elevated when the persistence-inducing trigger, in this case nutrient starvation, is applied longer. Interestingly, this high number of persister cells is not maintained but decreases again at later time points. We observed the same trend when cultures were treated with the aminoglycoside amikacin (Figure S1C).

#### Protein aggregation is a physiological change that occurs during stationary phase

Because of its previously reported connection to persistence, we assessed the dynamics of protein aggregation using quantitative image analysis (Figure 1C and D). Aggregates were identified either as phase-bright intracellular structures (further referred to as Ph aggregates) or as fluorescent IbpA-msfGFP foci (IbpA aggregates) (24). The chaperone IbpA is expressed upon protein aggregation and localizes to aggregates. Fluorescent fusions of IbpA can thus be used as a biosensor for protein aggregation, provided they do not trigger aggregation themselves (24, 25). Investigation of a wide array of different IbpA fusions showed that IbpA-msfGFP performed the best and can reliably identify protein aggregates (24). From Figure 1D it is clear that the number of cells with protein aggregates strongly increases during stationary phase and that Ph aggregates appear later than IbpA aggregates. These data suggest that protein aggregates develop gradually. Indeed, time lapse microscopy of single aggregates revealed that these structures are first recognized by IbpA-msfGFP and then gradually develop into dense phase-bright foci (Movie S1 and Figure S1D). By evaluating protein aggregation using IbpA-msfGFP and phase contrast microscopy, we can thus distinguish aggregates at an early stage of development (marked by IbpA-msfGFP) and a late stage of development (also visible in phase contrast microscopy).

#### The VBNC state and persistence are correlated to protein aggregation at the single-cell level

We next investigated whether there is a correlation between protein aggregation and bacterial dormancy manifested as persistence and the VBNC state. To assess the level of aggregation in persisters, we killed all antibiotic-sensitive cells with ofloxacin before seeding them onto agarose pads of fresh growth medium. Using time-lapse microscopy, we recorded for each cell whether or not it was able to resume growth, i.e. whether or not it is a persister, and its maximal fluorescence and phase contrast intensity at the start of the experiment. These values can be used to evaluate the level of IbpA and Ph aggregation, respectively (Figure S1E-F). This analysis showed that persisters have a much higher IbpA-msfGFP intensity, while Ph intensity increases only marginally (Figure 2A and B). Persisters therefore appear to be marked by early-stage protein aggregates. To assess the correlation between aggregation and the VBNC state, a similar strategy was followed but stationary-phase cells were directly seeded onto agarose pads without antibiotic treatment. Results indicate that cells unable to resume growth, i.e. VBNC cells, have a significantly higher Ph intensity (Figure 2C and D). Late-stage Ph aggregates are therefore associated with the VBNC state.

**Figure 2:**
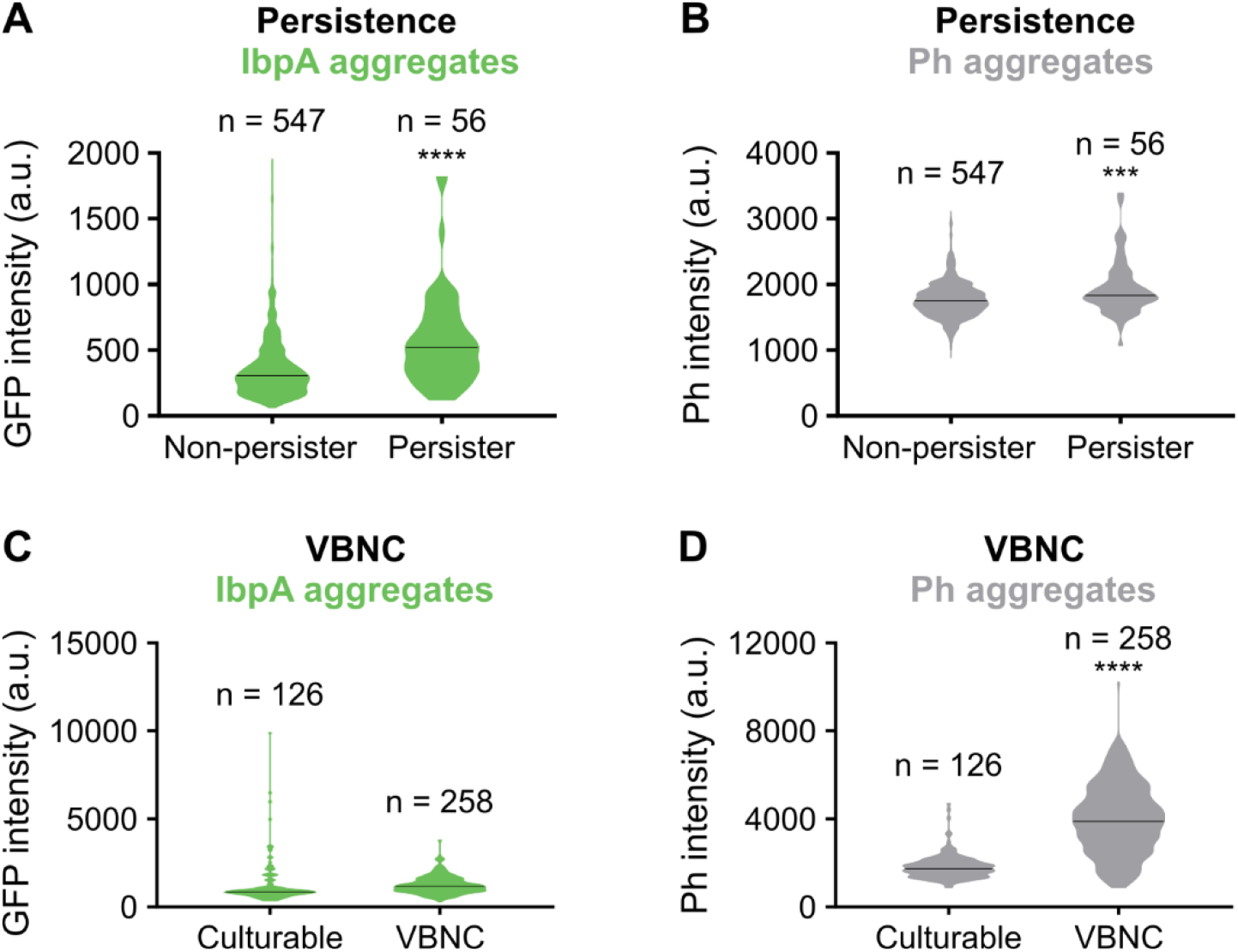
Persistence is correlated to early-stage IbpA-msfGFP aggregates while the deeply dormant VBNC state is correlated to late-stage Ph aggregates at the single-cell level. A-B) *E. coli ibpA-msfGFP* was grown for 24 hours and treated with ofloxacin to kill all antibiotic-sensitive cells. Regrowth of persisters on fresh medium was followed by time lapse microscopy. Cells were classified into two groups, persister or non-persister, based on whether or not they were able to resume cell division within 16 hours. Violin plots of the maximal IbpA-msfGFP fluorescence intensity (A) or maximal intensity in the phase contrast channel (B) are shown. C-D) To sample cells at different stages of dormancy, *E. coli ibpA-msfGFP* was grown for 24, 40 or 72 hours and regrowth on fresh medium was monitored by time lapse microscopy. Cells were classified into two groups, culturable or VBNC, based on whether or not they were able to resume cell division within 16 hours. Violin plots of the maximal IbpA-msfGFP fluorescence intensity (C) or maximal intensity in the phase contrast channel (D) are shown. Black lines in violin plots indicate median values. n is the number of cells measured. *** p < 0.001, **** p < 0.0001.

Since persister cells are marked by early-stage aggregates and deeply-dormant VBNC cells contain late-stage phase-bright aggregates, our results suggest that progressive protein aggregation drives dormancy development from the antibiotic-tolerant persister phenotype to the deeply-dormant VBNC state.

#### The persister protein ObgE induces aggregation

After establishing that persistence is associated with protein aggregation, we asked whether any of the proteins known to induce persistence do so by influencing aggregation. Overexpression of the GTPase *obgE*, which was shown to increase persister levels (26), indeed strongly accelerates protein aggregation (Figure 3A, B and C). Moreover, the aggregates that are formed are much larger and more intense (Figure 3A and Figure S2A-D). However, overexpression of *mCherry* from the same plasmid and promoter did not affect aggregation (Figure S2E and F), indicating that protein abundance *per se* is insufficient to trigger aggregation.

**Figure 3:**
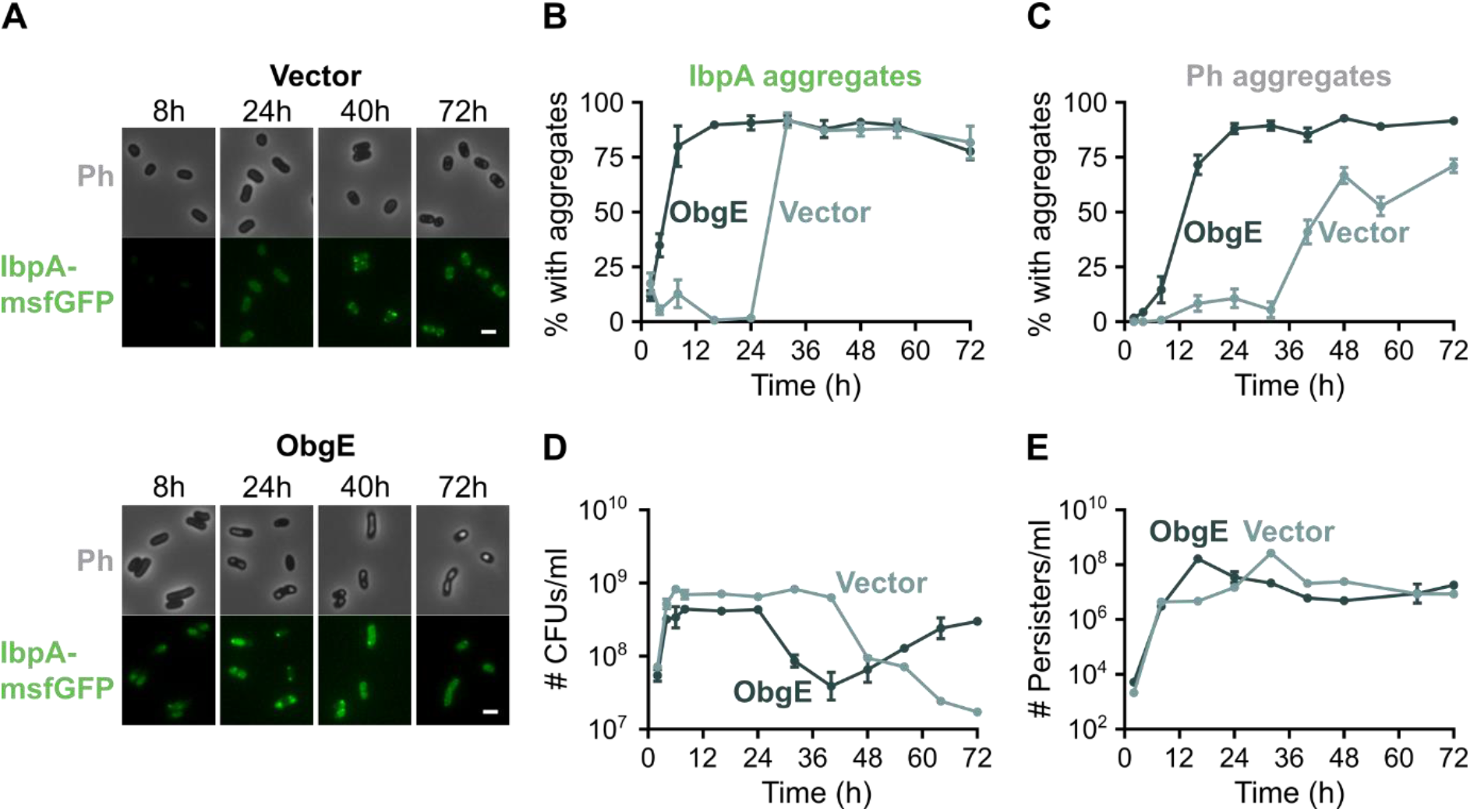
The persister protein ObgE accelerates protein aggregation, the development of persistence and entry into the VBNC state. *E. coliibpA-msfGFP* with pBAD33Gm (Vector) or pBAD33Gm-*obgE* (ObgE) was grown for 72 hours in the presence of the inducer arabinose. A) At different time points, microscopy pictures were taken. Scale bar, 2 μm. B-C) Quantitative analysis of microscopy images was performed to determine the percentage of cells that carry protein aggregates. Aggregation was evaluated by the presence of fluorescent IbpA-msfGFP foci (B) and phase-bright structures (C). For every repeat and every time point at least 50 cells were analyzed. D) At different time points, the number of CFUs/ml was determined. E) The absolute amount of persister cells after treatment with ofloxacin was followed over time. Data are represented as averages ± SEM, n ≥ 3. Ph, phase bright.

In accordance with the changes in protein aggregation, *obgE* overexpression also strongly accelerates the transition into the VBNC state (Figure 3D). Because the number of viable cells does not vary much through time (Figure S2G and H), the decrease in CFUs seen in Figure 3D is almost exclusively caused by cells that become VBNC. Intriguingly, in ObgE-induced samples culturability is restored again at late time points. This switch back to the culturable state is accompanied by the appearance of cells that lack protein aggregates (Figure S2I).

Additionally, persister development is also strongly accelerated by *obgE* overexpression (Figure 3E). Surprisingly, however, the maximal persister level obtained is the same regardless of whether or not *obgE* is overexpressed. It is simply reached sooner in the presence of excess ObgE. At the time points we usually assess persistence (16-24 hours), the persister level of the ObgE sample is higher, which explains why we previously reported that *obgE* expression increases persistence (26). However, based on the dynamic persister measurements performed here, we conclude that ObgE accelerates persister development rather than increasing the maximal persister level.

Importantly, also for the vector control and the ObgE sample, we could demonstrate that persisters show increased IbpA-msfGFP fluorescence, while VBNC cells have a higher Ph intensity (Figure S3). These data therefore further support our hypothesis that progressive protein aggregation drives the gradual development of dormancy.

After establishing that overexpression of *obgE* strongly accelerates the development of protein aggregates, we wondered whether this GTPase also affects aggregation at wild-type levels without overexpression. To test this, we used IbpA-msfGFP fluorescence as a proxy for protein aggregation and simultaneously measured the cellular concentration of ObgE by creating a genomic fusion with mCherry and measuring red fluorescence. Representative flow cytometry plots and corresponding Pearson’s R values show that there is a strong correlation between protein aggregation (IbpA-msfGFP) and ObgE levels (ObgE-mCherry), while no or only a weak correlation between red and green fluorescence could be detected in strains that carried only one fluorescent reporter (Figure 4A). Moreover, this difference in the strength of correlation was found to be reproducible (Figure 4B) and statistically significant in all repeats (p < 0.0001). Since cells that contain more ObgE show an increase in protein aggregation, we conclude that also at wild-type levels ObgE plays a role in triggering protein aggregation.

**Figure 4:**
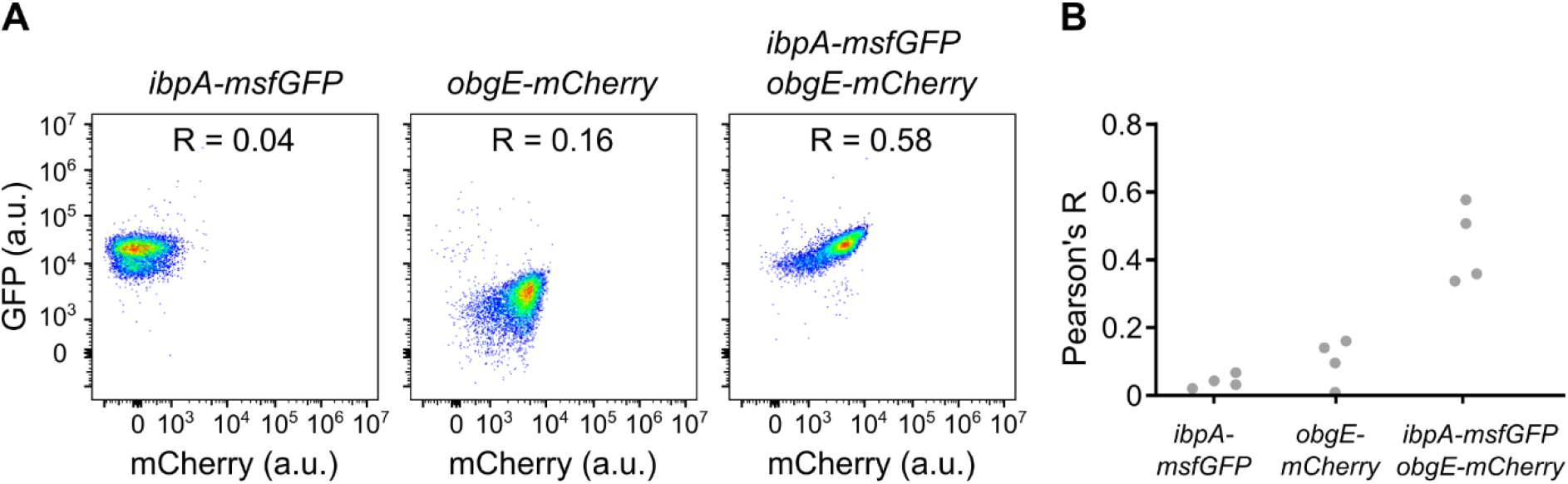
Wild-type levels of ObgE correlate to the amount of protein aggregation. The fluorescence of *E. coli* cells containing the genomic markers *ibpA-msfGFP, obgE-mCherry* or both were measured by flow cytometry after 24h of growth. Four different biological repeats were performed. A) Representative flow cytometry plots and corresponding Pearson’s correlation coefficients are shown. B) The correlation coefficients for all four repeats are plotted. Statistical analysis indicates that the Pearson’s R values obtained for the sample that contains both *ibpA-msfGFP* and *obgE-mCherry* are significantly higher in all repeats (p < 0.0001).

#### ObgE-induced aggregation requires entry into stationary phase

The most straightforward way in which an excess of ObgE could induce aggregation is by acting as a nucleation factor that initiates and drives aggregation through improper or incomplete folding (27). However, it is unlikely that ObgE acts as a nucleator. First, ObgE is not highly prone to aggregation since it is not commonly found in protein aggregates (14, 28, 29) and is highly soluble in cell-free translation systems (30, 31). Indeed, the aggregation propensity of ObgE, as measured by the Tango algorithm (32), is much lower than that of the known aggregation-prone protein MetA (33) (925.4 for ObgE versus 2641.8 for MetA). Second, *obgE* overexpression in itself is insufficient to trigger aggregation. Whereas MetA triggers aggregation already in exponential phase and also during balanced growth (where the transition into stationary phase is prevented by repeated dilution of a growing exponential-phase culture), ObgE is incapable of doing so (Figure 5A and S4). ObgE thus requires a stationary-phase signal to trigger aggregation. This dependency of ObgE-induced aggregation on specific growth conditions combined with the fact that ObgE is not highly prone to aggregation, leads us to conclude that ObgE does not act as a nucleation factor but triggers aggregation by a different mechanism that is activated in stationary phase.

**Figure 5:**
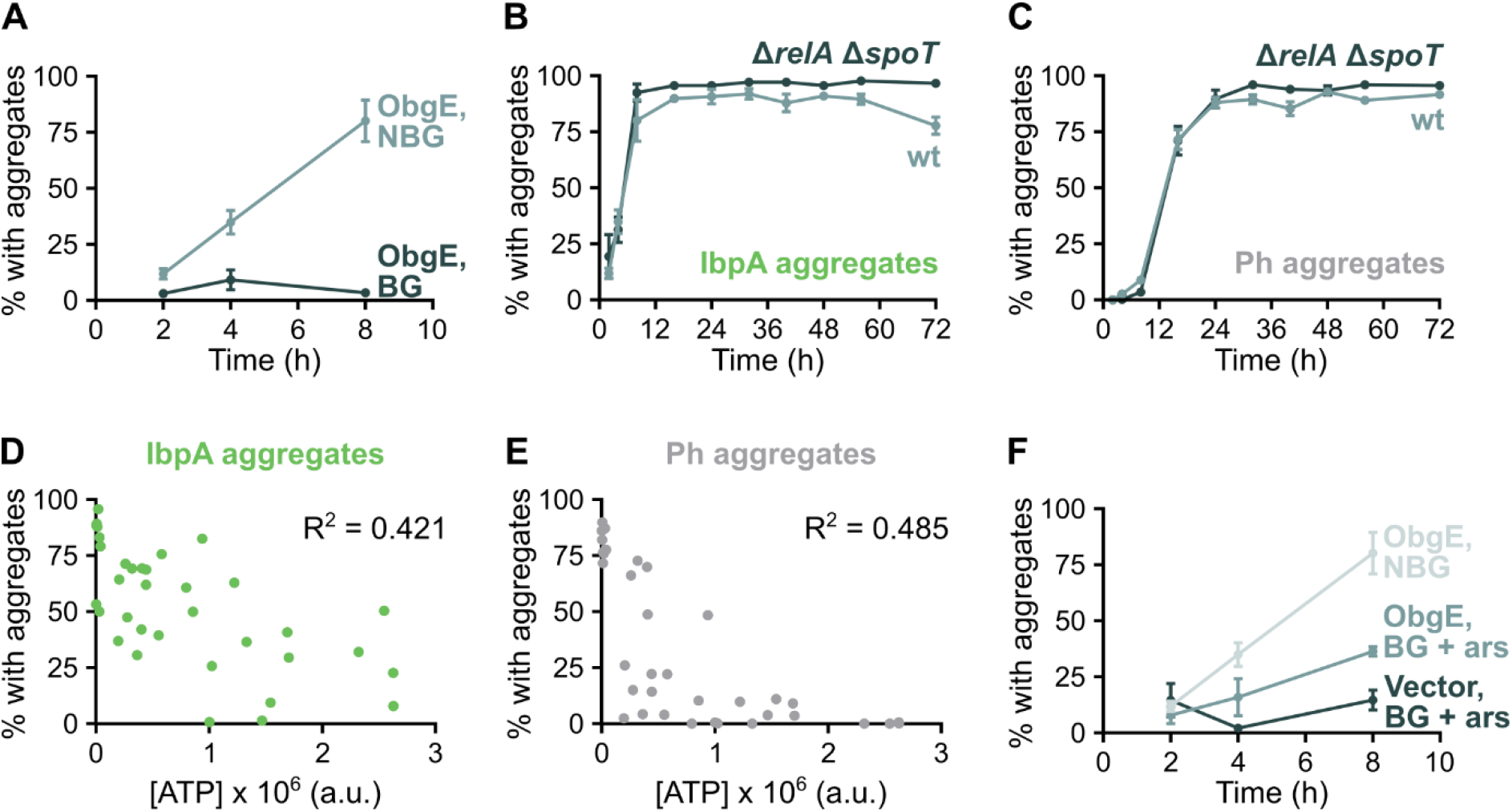
ATP depletion associated with entry into stationary phase is required for ObgE to induce protein aggregation. A) *E. coli* overexpressing *obgE* was grown normally (non-balanced growth, NBG) or maintained in balanced growth (BG). At different time points, the percentage of cells that carry IbpA aggregates was determined. B-C) The percentage of cells of a wild-type and *ΔrelA ΔspoT*culture that carry protein aggregates upon *obgE* overexpression was determined by quantitative analysis of microscopy images. Aggregation was evaluated by the presence of fluorescent IbpA-msfGFP foci (B) and phase-bright structures (C). D-E) There is a correlation between cellular ATP levels and the tendency to develop either IbpA (D) or Ph (E) protein aggregates. Both correlations are highly significant (p < 0.0001). F) *E. coli ibpA-msfGFP* with pBAD33Gm (Vector) or pBAD33Gm-*obgE* (ObgE) was maintained in balanced growth (BG) with 0.5 mM arsenate (ars) to lower ATP levels. At different time points, the percentage of cells that carry IbpA aggregates was determined. The ObgE NBG curve from (A) is also shown to allow easy comparison.

#### (p)ppGpp is not necessary for ObgE-induced protein aggregation

In search of the stationary-phase signal that is required by ObgE to trigger aggregation, we turned to the alarmone (p)ppGpp that is produced upon nutrient starvation (34). We previously showed that ObgE requires (p)ppGpp to induce persistence (26), making this molecule an interesting candidate. To our surprise, however, the amount of aggregation upon *obgE* overexpression is unaffected in a *ΔrelA ΔspoT* strain that is unable to produce (p)ppGpp (Figure 5B and C). (p)ppGpp is thus not the stationary-phase signal needed by ObgE to trigger aggregation.

The fact that obgE overexpression no longer induces persistence in the absence of (p)ppGpp (26) and that ObgE-induced protein aggregation proceeds normally in a *ΔrelA ΔspoT*strain seems to contradict the correlation between aggregation and persistence we report here. We therefore investigated persistence and dormancy upon *obgE* overexpression in the *ΔrelA ΔspoT* strain in more detail. Persister measurements indeed show that fewer cells are persistent in the absence of (p)ppGpp (Figure S5A), thereby confirming our previously published results (26). On the other hand, VBNC cells are present from very early on and are much more abundant in the *ΔrelA ΔspoT* strain (Figure S5B). These results suggest that the transition into the VBNC state occurs sooner and more frequently in this mutant. In line with this finding, we see that, in the absence of *obgE* overexpression, aggregation is strongly accelerated in the *ΔrelA ΔspoT* mutant (Figure S5C and D) and aggregates are much larger and more intense (Figure S5E-I). These data indicate that protein aggregation occurs more extensively in the absence of (p)ppGpp. We therefore hypothesize that the accelerated and more extreme protein aggregation in the *ΔrelA ΔspoT* strain pushes cells from the persister state into the deeply-dormant VBNC state more quickly, thereby explaining both the increased number of VBNC cells and the absence of an increased persister level.

#### ATP depletion is required for ObgE-induced protein aggregation

Another hallmark of entry into stationary phase is a decrease in cellular energy levels. Moreover, ATP is known to act as a hydrotrope that can prevent aggregation and increase protein solubility (35, 36). By modulating ATP levels through the addition of arsenate, we confirm that there is a highly significant (p < 0.0001), although rather weak, correlation between cellular ATP levels and protein aggregation at the population level (Figure 5D and E), as was also reported previously (14). Because of the connection between ATP concentration and aggregation, we assessed the role of ATP depletion in ObgE-induced aggregation by adding arsenate to cultures maintained in balanced growth. Low arsenate levels could partially restore ObgE’s ability to induce aggregation (Figure 5F), while having no effect on a control culture, hinting at an ObgE-specific and ATP-dependent aggregation mechanism. It therefore appears as though a lowered ATP concentration is the stationary-phase signal needed by ObgE to induce aggregation.

#### Protein aggregates are enriched in proteins involved in translation

Finally, we studied the composition of starvation- and ObgE-induced protein aggregates isolated at several different time points. First, using quantitative interpretation of Coomassie stained gels, we established that the amount of aggregated protein increases through time (Figure 6A and B), again highlighting the progressive nature of protein aggregation. Moreover, as expected based on their size, ObgE-induced aggregates have a higher total protein content. Second, we identified the aggregated proteins by mass spectrometry (MS) and detected around 500-600 proteins in each sample (Supplemental table 1). To gain a more high-level view of the types of proteins that aggregate, we performed enrichment analyses based on Clusters of Orthologous Group (COG) functional categories (Figure 6C and Figure S6A and B) (37). Strikingly, we found that four COG functional categories were significantly enriched in all time points for both the control and ObgE samples. These categories are C (energy production and conversion), F (nucleotide transport and metabolism), J (translation, ribosomal structure and biogenesis) and O (post-translational modification, protein turnover and chaperones), with category J being the most highly enriched. Proteins involved in these functions are thus more likely to aggregate than expected based on random chance. Interestingly, the level of enrichment between the control and ObgE samples is almost identical (Figure 6C and Figure S6A and B), indicating that the same types of proteins aggregate, regardless of whether aggregation is allowed to proceed naturally upon starvation or whether it is accelerated by *obgE* overexpression.

**Figure 6:**
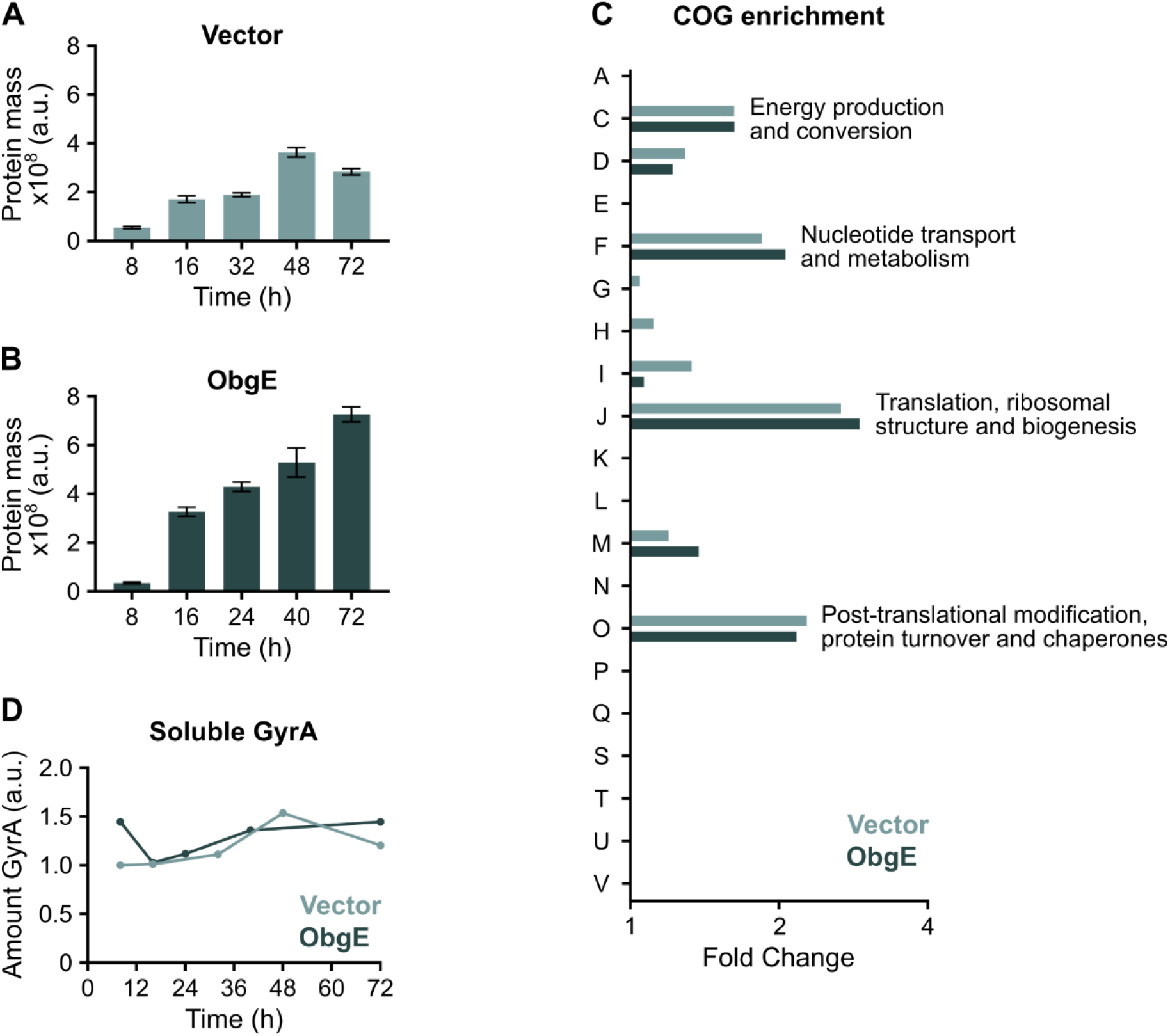
Biochemical analysis of protein aggregates reveals their time-dependent development and non-random composition. A-B) After isolation of protein aggregates from *E. coli*carrying an empty expression vector (A) or overexpressing *obgE* (B) at several different time points, Coomassie staining on protein gels was performed. Quantitative interpretation of these Coomassie stained gels is shown as ‘protein mass’. Data are represented as averages ± SEM, n = 5. C) COG functional categories were assigned to all aggregated proteins identified by MS and an enrichment analysis was performed. The amount of enrichment of all identified categories at the time point where Ph aggregation reaches its plateau is shown (Vector = 48h, ObgE = 24h). Significantly enriched COG categories (C, F, J, O; p < 0.05) are highlighted by including the description of the category inside the figure. All descriptions; A: RNA processing and modification, C: Energy production and conversion, D: Cell cycle control, cell division, chromosome partitioning, E: Amino acid transport and metabolism, F: Nucleotide transport and metabolism, G: Carbohydrate transport and metabolism, H: Coenzyme transport and metabolism, I: Lipid transport and metabolism, J: Translation, ribosomal structure and biogenesis, K: Transcription, L: Replication, recombination and repair, M: Cell wall/membrane/envelope biogenesis, N: Cell motility, O: Post-translational modification, protein turnover and chaperones, P: Inorganic ion transport and metabolism, Q: Secondary metabolites biosynthesis, transport and catabolism, S: Function unknown, T: Signal transduction mechanisms, U: Intracellular trafficking, secretion and vesicular transport, V: Defense mechanisms. D) The amount of GyrA detected in the soluble protein fraction by western blot with anti-GyrA antibodies is shown. The detected signals were normalized to the first time point of the vector control (8h).

Despite the highly similar functional composition of starvation- and ObgE-induced protein aggregates, there are differences when looking at the individual protein level (Supplemental table 2). A total of 131 proteins are detected in significantly higher amounts when *obgE* is overexpressed. A COG enrichment analysis of these 131 proteins reveals that only category, J (translation, ribosomal structure and biogenesis), is significantly enriched (p value = 7.59 x 10^-6^, fold change = 3.24). This means that, in comparison to starvation-induced aggregation, ObgE causes mainly proteins involved in translation to aggregate more frequently. Since ObgE is known to interact with the 50S ribosomal subunit(38), we hypothesize that this interaction is responsible for the observed increase in aggregation of translation-related proteins. Indeed, many 50S ribosomal proteins are more abundantly present in ObgE-induced aggregates (Supplemental table 2). If true, the widely conserved interaction of ObgE with the ribosome is important for the regulation of bacterial dormancy and survival of stressful conditions such as antibiotic insults.

Finally, we asked if and how aggregate composition could explain persister development. The antibiotic-tolerant persister state is classically attributed to the lack of activity of the antibiotic target (7), although this is not universally true (6). Since GyrA, the main target of ofloxacin in *E. coli* (39), is detected in aggregates (Supplemental table 1), we wondered whether sequestration and subsequent inactivation of cytoplasmic GyrA in protein aggregates is responsible for persister development. We therefore measured the remaining amount of soluble GyrA and established that it remains more or less constant through time (Figure 6D), even though the persister level changes. Persistence is therefore not obtained through sequestration of GyrA in protein aggregates, which is in accordance with the finding that persisters do experience DNA damage upon ofloxacin treatment and that the antibiotic target is thus not completely inactivated (6). Rather, we believe that aggregation leads to persister development through the aggregation of a wide variety of important and essential proteins resulting in cellular dormancy.

## Discussion

Based on our results, we propose the following model to explain the development of both persistence and the VBNC state upon starvation in stationary phase (Figure 7). When nutrients become scarce, ATP levels decrease and proteostasis cannot be maintained. As a result, protein aggregates start to develop. Aggregation occurs progressively and different stages of protein aggregation are linked to different dormancy depths. On average, persisters are cells that contain aggregates in an early stage of development, while VBNC cells carry more developed aggregates. These findings lead us to hypothesize that early-stage aggregates render bacteria sufficiently dormant to survive antibiotic treatment, yet active enough to resume growth when provided with fresh nutrients. Late-stage protein aggregates, on the other hand, induce a deeper state of dormancy, i.e. the VBNC state, from which cells cannot readily wake up even under favorable conditions that normally would support growth.

**Figure 7:**
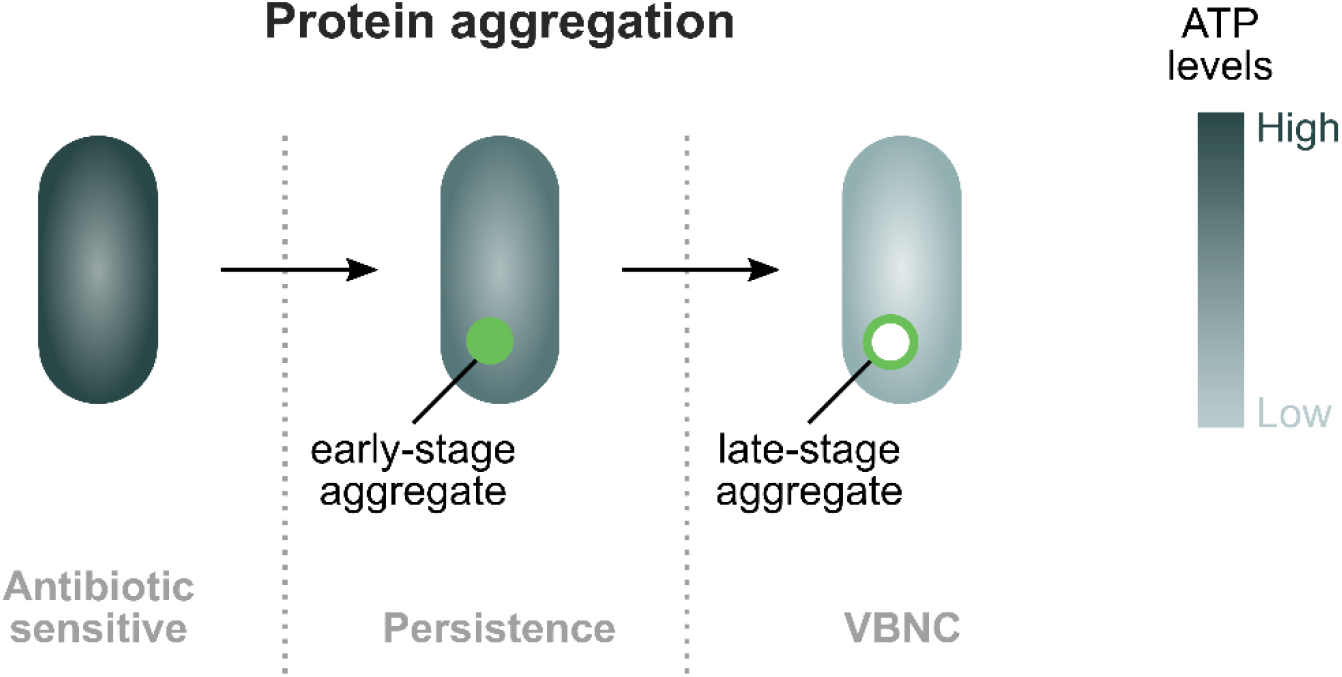
Model for the development of bacterial dormancy through protein aggregation. Upon nutrient starvation in stationary phase, cellular ATP concentrations decrease. As a consequence, proteostasis cannot be maintained and protein aggregates develop. Protein aggregation occurs progressively. In an early stage of development, protein aggregates are marked by the small chaperone IbpA. Later on, they further develop into phase-bright structures. These different stages of protein aggregation are tightly coupled to different stages of bacterial dormancy. Persister cells are characterized by early-stage IbpA aggregates, while deeply-dormant VBNC cells contain more developed aggregates that can be seen in phase contrast microscopy.

However, it is important to note that not all cells that carry early-stage aggregates are persisters and not all cells with late-stage aggregates are in the VBNC state. We therefore hypothesize that it is not the mere presence of these aggregates that results in the associated dormancy phenotype. Rather, we suspect that the level of dormancy is determined by the specific aggregate composition. Since proteins involved in translation, ribosomal structure and biogenesis are most highly enriched in aggregates, we suspect that translation plays a pivotal role in protein aggregation and subsequent dormancy development. Supporting this hypothesis is the finding that directly inhibiting protein expression increases bacterial persistence dramatically (40, 41). Moreover, we find that overexpression of *obgE* strongly accelerates the development of protein aggregates, persistence and the transition into the VBNC state and simultaneously increases the amount of aggregated translation-related proteins. Since ObgE is known to interact with the ribosome (38) it is possible that ObgE modulates aggregation and dormancy through this ribosomal interaction. Importantly, ObgE’s effect on protein aggregation is not a mere artefact of protein overexpression since we here demonstrate that, also at wild-type levels, the amount of ObgE is correlated to the level of protein aggregation.

Since we show that different stages of protein aggregation are associated with different dormancy depths, i.e. persistence and the VBNC state, and that protein aggregation occurs progressively, our findings provide very strong experimental support for the hypothesis that persisters and VBNC cells represent different stages of the same developmental dormancy program that is driven by – or at least associated with – protein aggregation. Additionally, it is important to note that a cell is considered to be a persister only when it can resume growth after antibiotic treatment (1). Changed persister levels should therefore be interpreted with care. A decrease in persistence can be due to less cells surviving antibiotic treatment, less cells being able to resume growth (i.e. more VBNC cells), or a combination of both. Because persisters can develop into VBNC cells, the time at which persistence is assessed will strongly influence the outcome. Indeed, whereas it is well known that persister fractions rise upon entry into stationary phase (22), we here show that persister levels are also highly dynamic throughout stationary phase. A previous study likewise reported that the detected persister fraction depends on the age of the inoculum used (42). Moreover, we show that the timing of persister development can be influenced, for example by the known persister protein ObgE.

Taken together, our data indicate that, contrary to current practice, the dormant phenotypes persistence and the VBNC state should be studied within the same conceptual framework, at least when induced by starvation. Especially considering that some VBNC cells can resume growth *in vivo* (43–46), there is no solid rationale to limit research on antibiotic tolerance to persistence and exclude VBNC cells that represent the next stage in dormancy development. We therefore argue that, to fully understand the influence of specific conditions on bacterial dormancy, persister levels should be followed through time and the number of VBNC cells should be determined simultaneously.

## Materials and methods

### Bacterial strains and growth conditions

Experiments were performed with *E. coli* BW25113 (47). In all experiments where aggregation was quantified, *E. coli* BW25113 *ibpA-msfGFP* was used. The *ibpA-msfGFP* allele was introduced using P1 phage transduction starting from donor strain *E. coli* MG1655 *ibpA-msfGFP* (24). A genomic *obgE-mCherry* fusion was created by inserting *mCherry-FRT-Km^R^-FRT* into the genome downstream of *obgE* by homologous recombination, thereby eliminating the *obgE* stop codon. The DNA fragment needed for homologous recombination was obtained through SOEing PCR using primers targeting homologous genomic regions (SPI13305 & SPI13306 and SPI13309 & SPI13310) and primers that amplify *obgE-mCherry-FRT-Km^R^-FRT* present in *pBAD33-obgE-mCherry-FRT-Km^R^-FRT* (lab collection) (SPI13307 & SPI13308, see Supplemental table 3 for primer sequences). After incorporation, the FRT-flanked kanamycin resistance cassette was removed from the genome (47). *E. coli* BW25113 *ΔrelA ΔspoT* (48) was used to interrogate the effect of (p)ppGpp on aggregation.

Where applicable, *obgE* was overexpressed from plasmid pBAD33Gm-*obgE* (49). The corresponding *E. coli* strain carrying the empty pBAD33Gm vector (49) was used as a control. pBAD33Gm-*mCherry* was constructed by amplification of *mCherry* from pBAD/*Myc*-His *A-mCherry* (26) with primers SPI12788 and SPI12789 (see Supplemental table 3 for primer sequences). The *mCherry* amplicon was inserted into pBAD33Gm by digestion with SacI and HindIII and subsequent ligation. In all cases, expression from pBAD33Gm plasmids was induced by adding 0.2 % w/v arabinose. MetA was overexpressed from pCA24N (50) by the addition of 100 μM IPTG.

For all tests, overnight cultures were diluted 100 times in lysogeny broth (LB) containing the appropriate antibiotics (chloramphenicol 35 μg/ml, gentamicin 25 μg/ml, kanamycin 40 μg/ml, spectinomycin 50 μg/ml) and incubated at 37°C with continuous shaking at 200 rpm. At indicated time points after dilution, assays were performed.

### Determining the amount of culturable and viable but nonculturable cells

The amount of culturable cells was determined by measuring the number of CFUs per ml. Serial dilutions were prepared in 10 mM MgSO4 and plated on LB medium containing 1.5 % agar. After overnight incubation at 37°C, colonies were counted and the number of CFUs per ml was calculated.

To determine the number of VBNC cells, the number of CFUs per ml was compared to the amount of viable cells per ml. The latter was determined in the following way. First, the total number of cells (alive or dead) per ml was determined using Thermo Fisher’s Bacteria Counting Kit for flow cytometry. Briefly, approximately 10^6^ bacteria were stained with SYTO^™^ BC following manufacturer’s instructions. A known amount of fluorescent microsphere beads was added to the stained cell preparation. These microspheres can easily be distinguished from bacteria and were used as a standard to accurately determine the total number of cells in the original culture. Analyses were performed using a BD Influx cell sorter equipped with 488 nm and 561 nm lasers and standard filter sets. Second, the percentage of viable cells in the population was determined by staining cells with SYTOX Green, a green fluorescent dye that can only enter cells that have lost membrane integrity and are therefore considered dead. Cultures were diluted 100 times in PBS containing 0.5 μM SYTOX Green. Samples were incubated for 15 min in the dark at room temperature and analyzed using a BD Influx cell sorter. To calculate the concentration of viable cells in the population, the total number of cells per ml determined in the first step was multiplied by the percentage of non-fluorescent – and therefore viable – cells in the population.

Additionally, viability of cultures after 72 hours of incubation was confirmed by staining with the redox-sensitive dye RedoxSensor Green according to manufacturer’s instructions. As a negative control, *E. coli* was treated with 10 mM sodium azide as recommended by the manufacturer. Cell vitality was assessed by flow cytometry using a BD Influx cell sorter.

### Persister assays

To determine the amount of persister cells, an aliquot of the culture was taken at appropriate time points and treated with 5 μg/ml ofloxacin for 5 hours. Afterwards, cells were washed twice with 10 mM MgSO4. Serial dilutions were made in 10 mM MgSO4 and plated on non-selective LB medium containing 1.5 % agar. After 48 hours of incubation at 37°C, CFUs were determined.

### Microscopy analyses

All microscopy analyses were performed using a Nikon Ti-E inverted microscope equipped with Qi2 CMOS camera and temperature controlled cage incubator. For snapshot analyses performed at discrete time points, cells were spotted onto pads of 10 mM MgSO4 with 2 % w/v agarose. For time lapse analyses, agarose pads of LB medium were used and cells were grown at 37°C. To assess the correlation between protein aggregation and the VBNC state (Figure 2A and B), stationary-phase cultures were spotted onto agarose pads of fresh LB medium after 24, 40 or 72 hours of incubation and growth was followed for 16 hours. To assess the correlation between aggregation and persistence (Figure 2C and D), stationary-phase cultures were treated with 5 μg/ml ofloxacin for 5 hours after 24 hours of incubation. Cells were washed twice in 10 mM MgSO4 and spotted onto agarose pads of fresh LB medium. Growth was followed for 16 hours. To follow the development of protein aggregates (results presented in Figure S1C), exponential-phase cells overexpressing *obgE* were spotted on agarose pads of spent LB medium, thereby limiting the number of cell divisions before aggregation occurs. Spent LB medium was generated by growing *E. coli* cells for 24 hours, centrifuging at 4000 rpm for 20 min and filtering the supernatant using a 0.22 μm pore size. This experiment was performed with *obgE* overexpression because we were unable to track full aggregate development in a wild-type strain. Using the wildtype, the detected IbpA-msfGFP foci never developed into late-stage Ph aggregates on solid medium.

For all experiments, image analysis was performed using MicrobeJ (51). This ImageJ plug-in was used to determine the presence, intensity and size of IbpA-msfGFP fluorescent foci and phase-bright structures. For time lapse analyses correlating protein aggregation to bacterial dormancy, IbpA-msfGFP and Ph intensities were normalized to the background in order to allow comparison of intensities across different experiments with slightly different light settings. Because corrected Ph intensities often have negative values (cells are darker than the background), which may be confusing for the interpretation of results, all corrected Ph intensities were transformed (+ 2000 a.u.) to obtain positive values.

### Determining the correlation between wild-type ObgE levels and protein aggregation

Flow cytometry experiments were performed on a CytoFLEX S instrument (Beckman Coulter Life Sciences) equipped with 405 nm, 488 nm and 561 nm lasers. Cells were grown for 24h and diluted 1000x in PBS before measuring. Bacterial cells were gated based on FSC and SSC values and for each sample 10.000 cells were collected. Blead-through of fluorophores in other channels was corrected by applying a compensation matrix to the measured fluorescence values following manufacturer’s instructions. Pearson’s R values were calculated for log-transformed and compensated fluorescence values.

### Aggregation predictions using Tango

The Tango algorithm (32) was applied using the following settings; Nterm N, Cterm N, pH 7.5, Temperature 310.15, Ionic strength 0.1, Sequence 1. For respectively ObgE and MetA, the amino acid sequences with GenBank identifiers VWQ04101.1 and SYX52533.1 were used.

### Balanced growth

To maintain balanced growth, cultures were diluted 3- or 4-fold every time the OD 595 nm reached values between 0.25 and 0.3. Dilutions were made in pre-warmed LB medium with the appropriate antibiotics and additives (0.2 % w/v arabinose, 100 μM IPTG and/or 0.5 mM arsenate).

### ATP measurements

ATP measurements were performed using the BacTiter-Glo^™^ Microbial Cell Viability Assay according to manufacturer’s instructions. Briefly, cells were washed to remove any potential ATP contamination present in the growth medium. Equal volumes of the cell suspension and the BacTiter-Glo reagent were mixed and incubated for 5 min before measuring luminescence using a Synergy Mx Monochromator-Based Multi-Mode Microplate Reader (BioTek). Signal intensities were corrected for cell numbers by dividing the luminescence signal by the OD 595 nm of the cell suspension. ATP levels were modulated by adding different concentrations of arsenate (0.1, 1 and 10 mM) to bacterial cultures and measuring at different time points, and hence in different growth phases of the culture.

### Isolation of protein aggregates and soluble protein fractions

Protein aggregates were isolated at several different time points as described previously (28). The time points chosen reflect different developmental stages of protein aggregates and cellular dormancy; aggregates were isolated when the plateau in aggregation (as detected by IbpA-msfGFP or Ph) was reached, when the maximal persister level was obtained and when the highest amount of VBNC cells were detected. For the vector control these correspond to 16, 32, 48 and 72 hours of incubation. For *obgE* overexpression, cultures were incubated for 8, 16, 24, 40 or 72 hours. At the corresponding time points, 25 ml of bacterial cultures was harvested by centrifugation at 4000 g for 30 min. For every condition and every time point, 5 repeats were performed. Cells were washed with 10 ml buffer A (50 mM HEPES, pH 7.5, 300 mM NaCl, 5 mM β- mercaptoethanol, 1.0 mM EDTA) and centrifuged at 4000 g for 30 min at 4°C. Pellets were dissolved in 20 ml buffer B (buffer A plus 1 μg/mL leupeptin, 0.1 mg/mL AEBSF (4-(2-aminoethyl)benzenesulfonyl fluoride hydrochloride)). Cells were lysed by a Glen Creston Cell Homogenizer with a pressure of 20 000 – 25 000 psi and by sonication with a Branson Digital sonifier (50/60 HZ) on ice for 2 min (cycles consist of 15 sec pulses at 50 % power with 30 sec pauses on ice). The lysed cell suspension was centrifuged at 11 000 g for 30 min at 4°C and the precipitated fraction was resuspended in 10 ml buffer D (buffer A with 0.8% (V/V) Triton X-100 and 0.1 % sodium deoxycholate). This step was repeated three times. Finally, aggregates were solubilized in 1 ml buffer F (50 mM HEPES, pH 7.5, 8.0 M urea).

Soluble protein fractions were separated by collecting 100 ml of bacterial cultures at the appropriate time points by centrifugation at 4500 g for 5 min at 4°C. The pellet was stored overnight at −20°C. Pellets were dissolved in PBS containing 1x EDTA-free protease inhibitor and benzonase and cells were lysed by sonication for 3 min (cycles consist of 30 sec pulses at 35% amplitude with 40 sec pauses on ice). Cell debris was removed by centrifugation at 4500 g for 5 min. Soluble and insoluble fractions were separated by high-speed centrifugation at 250 000 g for 20 min at 4°C.

### Coomassie staining

Isolated aggregates were loaded on SDS gels (4-15% Mini-PROTEAN^®^ TGX^™^ Precast Protein Gels, 10-well, 50 μl well volume) and stained by Coomassie blue (R250). Band quantification of the proteins on SDS gel was performed using Image Lab Software Version 6.1.

### Western blot analysis

At the indicated time points, soluble protein fractions were obtained as described above. NuPAGE LDS Sample Buffer (Invitrogen) and NuPage Sample reducing agent (Invitrogen) were added and samples were loaded on Novex 8% Tris-Glycine Gels (Invitrogen) in a Novex Bolt Mini Gel tank with MOPS buffer. Proteins were transferred onto a polyvinylidene fluoride (PVDF) membrane using preassembled Trans-Blot Turbo Transfer Packs and a Trans-Blot Turbo Transfer System (Bio-Rad). The voltage was set to 15 V and samples were run for 7 min. The membrane was washed with TBS supplemented with 0.1% Tween20 and blocked by incubation in blocking buffer (10% skimmed milk powder in TBS-Tween20) for 2 hours. Afterwards the membrane was washed with TBS-Tween20 and GyrA was detected with polyclonal anti-GyrA (1/200). After overnight incubation with the primary antibody, membranes were washed with TBS-Tween20. The secondary antibody, anti-rabbit IgG fused to horse radish peroxidase (1/50000), was added and the membrane was incubated for 55 minutes. After washing with TBS-Tween20, detection was done by adding ClarityTM Western ECL Substrate (Bio-Rad) and visualization with the Fusion FX imaging system (Vilber Lourmat). Pictures were taken and band intensities were quantified using ImageJ.

### MS analysis

MS analysis was performed according to a previously published protocol(28). Dithiothreitol (DTT) was added to purified aggregates to a total concentration of 0.02 M. Samples were incubated for 15 min at room temperature before IAA was added at a concentration of 0.05 M. Samples were incubated for 30 min protected from light. ABC was added at a final concentration of 0.11 M and 0.2 μg trypsin per 20 μg protein was added as well. Trypsin digestion was allowed to proceed for at least 16 h at 37°C. Peptides were then cleaned using C18 spin columns (Thermo Fisher Scientific) collected with elution buffer. The total peptides were then diluted in 5% CAN + 0.1% FA (10x) for injection in the Q Exactive Orbitrap mass spectrometer (Thermo Fisher Scientific). Five μl of each sample was loaded for UPLC separation into an Ultimate 3000 UPLC System (Dionex, Thermo Fisher Scientific), using an Acclaim PepMap100 pre-column (C18, 3 μm-100 Å) and a PepMap RSLC (C18, 2 μm, 50 μm-15 cm) both from Thermo Fisher Scientific. A linear gradient (300 μl/min) of buffer B (80% ACN, 0.08% FA) was applied: 0-4% for 3 min, 4-10% for 12 min, 10-35% for 20 min, 35-65% for 5 min, 65-95% for 1 min, 95% for 10 min, 95-5% for 1 min, and 5% for 10 min. The Q Exactive Orbitrap mass spectrometer (Thermo Fisher Scientific) was used for MS analysis. It was operated in positive ion mode and a nano spray voltage of 1.5 kV was applied with a source temperature of 250°C. For external calibration, Proteo Mass LTQ/FT-Hybrid ESI Pos Mode Cal Mix (MS CAL5-1EASUPELCO, Sigma-Aldrich) was used. Internal calibration was performed with the lock mass 445.12003. The instrument was operated in data-dependent acquisition mode with a survey MS scan at a resolution of 70 000 (fw hm at m/z 200) for the mass range of m/z 400-1600 for precursor ions, followed by MS/MS scans of the top 10 most intense peaks with +2, +3, +4 and +5 charged ions above a threshold ion count of 16 000 at 17 500 resolution using normalized collision energy of 25 eV with an isolation window of m/z 3.0 and dynamic exclusion of 10 sec. The Xcalibur 3.0.63 (Thermo Fisher Scientific) software was used for data acquisition. For protein identification, all raw data were converted using Proteome Discover 1.4 (Thermo Fisher Scientific) into mgf files and matched against the uniprot *E. coli* database using MASCOT 2.2.06 (Matrix Science). Parameters used are: parent tolerance of 10 ppm, fragment tolerance of 0.02 Da, variable modification oxidation of M, fixed modification with carbamidomethyl C, and up to one missed trypsin cleavage site. Results were imported into Scaffold 3.6.3. Proteins identified with 99% confidence that contain at least two identified peptides with a confidence level of over 95% were retained.

### Statistical analysis

Statistical analyses, except those presented in Figure 4, Figure 6, Figure S6 and Supplemental table 2, were performed using GraphPad Prism 8. Where applicable, two samples were compared using an unpaired t test. The statistics that are displayed in each figure, together with the number of repeats performed, are indicated in the figure legends. In general, data usually represent the mean ± SEM of at least 3 repeats. Various levels of significance were defined; * p < 0.05, ** p < 0.01, *** p < 0.001, **** p < 0.00001.

Pearson’s correlation coefficients were compared by first performing Fisher R-to-z transformations and subsequently evaluating the significance level of the obtained z values. For every repeat separately R coefficients of samples with only one fluorescent marker were compared to the R coefficient of the sample that contained both *ibpA-msfGFP* and *obgE-mCherry*. This generated four different z and p values for every comparison (one for every biological repeat). All of the comparisons revealed highly significant differences (p < 0.0001).

To identify aggregated proteins that are present in significantly different amounts in control and ObgE samples (Supplemental table 2), Welch’s t tests were performed and obtained p values were subjected to an FDR correction. The cut-off value for significance was set at p < 0.05.

COG functional categories were assigned to proteins identified inside aggregates using the eggNOG tool(52, 53). Enriched functional categories were identified by performing one-sided Fisher’s exact tests and applying a Bonferroni correction to the obtained p values. The cut-off value for significance was set at p < 0.05.

## Acknowledgements

We thank Abram Aertsen (KU Leuven) for the *E. coli* MG1655 *ibpA-msfGFP* strain. We are also thankful to SyBioMa, the Proteomics Core of the Biomedical Sciences Group of KU Leuven for their help and support in the analysis of aggregate composition. This research was funded by the KU Leuven Research Council (C16/17/006), FWO (G047112N; G0B2515N; G055517N), the Francqui Research Foundation, and the Flemish Institute for Biotechnology (VIB). The Switch Laboratory was supported by grants from VIB, the University of Leuven (“Industrieel Onderzoeksfonds”), the Funds for Scientific Research Flanders (FWO) and the Flemish Agency for Work and Innovation (VLAIO). LD, CB and EL received a fellowship from the FWO. DW received a fellowship from KU Leuven. PH received a fellowship from IWT.

## Author contributions

Conceptualization: LD, JM; Methodology; LD, JM; Investigation: LD, CB, DW, EL, PH, PM, LK, LK; Writing – Original draft: LD; Writing – Review & Editing: LD, CB, DW, LK, LK, FR, JS, JM; Visualization: LD; Supervision: JM.

## Supplemental information

**Figure S1:**
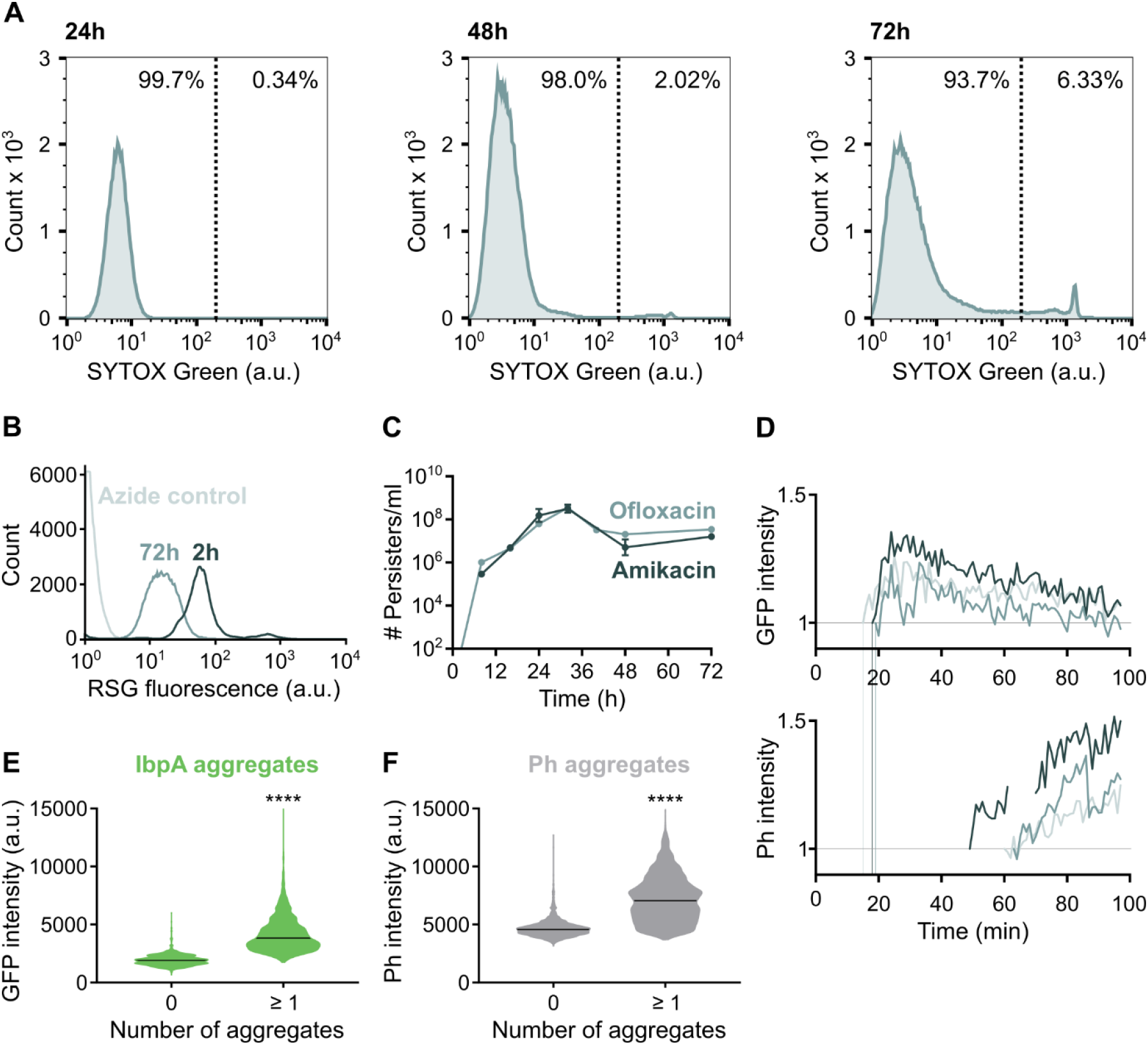
A) Viability of *E. coli*after long-term incubation in stationary phase is measured by staining cells at several different time points with the membrane-impermeable SYTOX Green dye. B) Viability of *E. coli* after long-term incubation in stationary phase is confirmed by staining cells that were grown for 72 hours with RedoxSensor Green (RSG). As a positive control for viability, exponential-phase cells that were incubated for 2 hours were used. As a negative control, cells were treated with sodium azide. C) Persistence of *E. coli* upon treatment with the fluoroquinolone ofloxacin or the aminoglycoside amikacin follow the same dynamics. Data are represented as averages ± SEM, n ≥ 3. D) The IbpA-msfGFP and phase contrast intensity of three single aggregates formed upon *obgE* overexpression was followed through time. To be able to track full development of protein aggregates on solid medium strong aggregation-inducing conditions were chosen; exponential-phase cells overexpressing *obgE* were spotted on agarose pads of spent LB medium, thereby limiting the number of cell divisions before aggregation occurs. E-F) Cellular IbpA-msfGFP fluorescence intensity and phase contrast intensity are correlated to the presence of protein aggregates identified as IbpA-msfGFP foci and phase-bright structures, respectively. After growing *E. coli ibpA-msfGFP* for 24, 40 or 72 hours, the maximal GFP and phase contrast intensity of each cell was recorded. Cells were divided into two categories based on whether they did or did not contain at least one fluorescent IbpA-msfGFP focus (E) or phase-bright structure (F). Black lines indicate median values. Over 2500 cells were analyzed. **** p < 0.0001.

**Figure S2:**
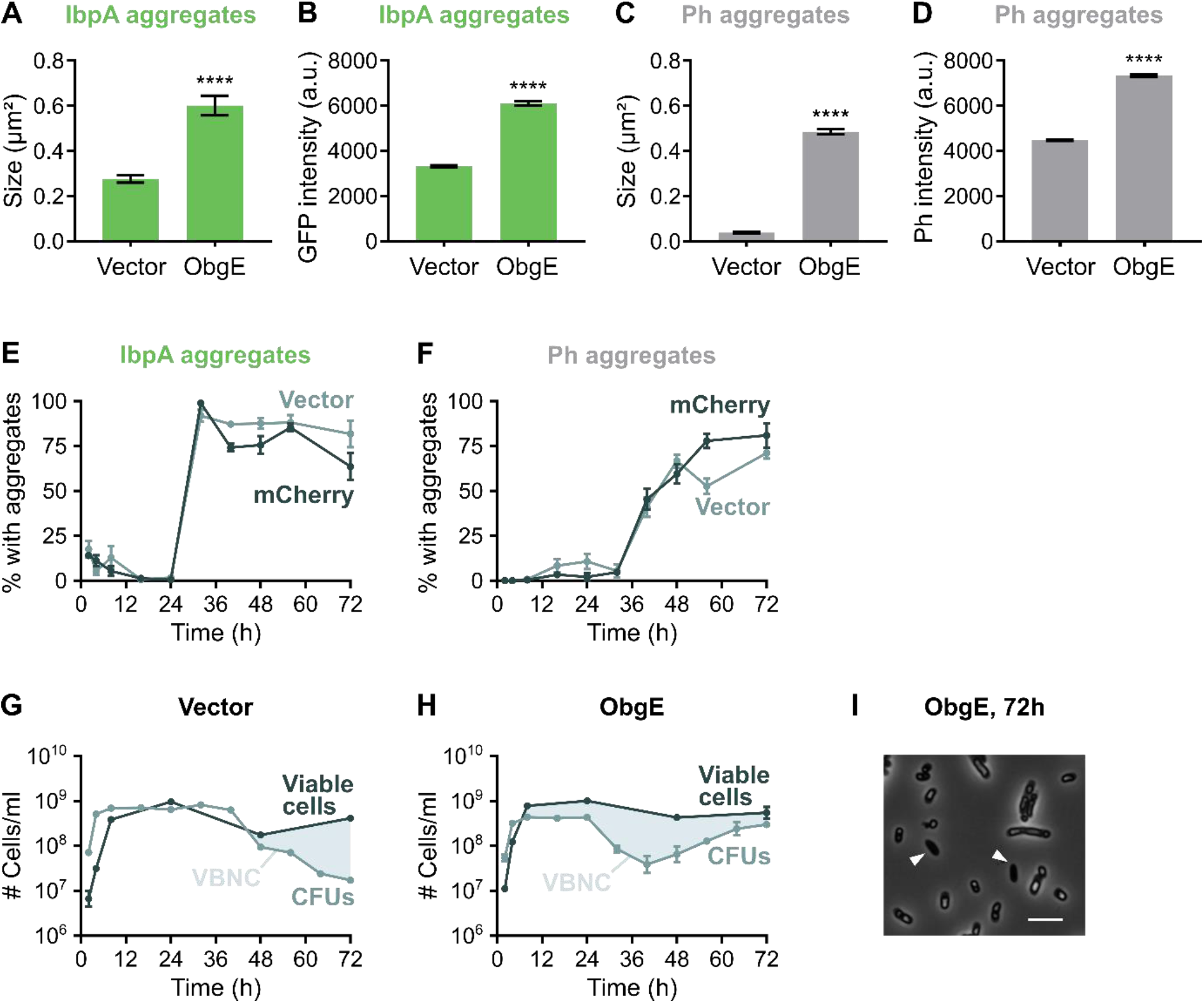
A-D) Protein aggregates that develop upon *obgE* overexpression are much larger and more intense than for the Vector control. This difference is demonstrated by quantitative measurements at time point 72h of the size (A) and intensity (B) of fluorescent IbpA-msfGFP foci and the size (C) and intensity (D) of phase-bright structures. Data are represented as averages ± SEM, n (number of cells analyzed) > 450, **** p < 0.0001. E-F) Overexpression of mCherry from the same plasmid and promoter as *obgE* does not induce aggregation. Quantitative analysis of microscopy images was performed to determine the percentage of cells that carry protein aggregates. Aggregation was evaluated by the presence of fluorescent IbpA-msfGFP foci (E) and phase-bright structures (F). For every repeat and every time point at least 50 cells were analyzed. Data are represented as averages ± SEM, n ≥ 3. G-H) A comparison of the number of CFUs/ml and the number of viable cells of *E. coli* pBAD33Gm (G) and *E. coli* pBAD33Gm-*obgE* (H) reveals the presence of VBNC cells in stationary phase. Data are represented as averages ± SEM, n ≥ 3. I) Microscopy image of *E. coli* overexpressing *obgE*for 72 h. Arrowheads point to cells that do not contain a protein aggregate. Scale bar, 5 μm.

**Figure S3:**
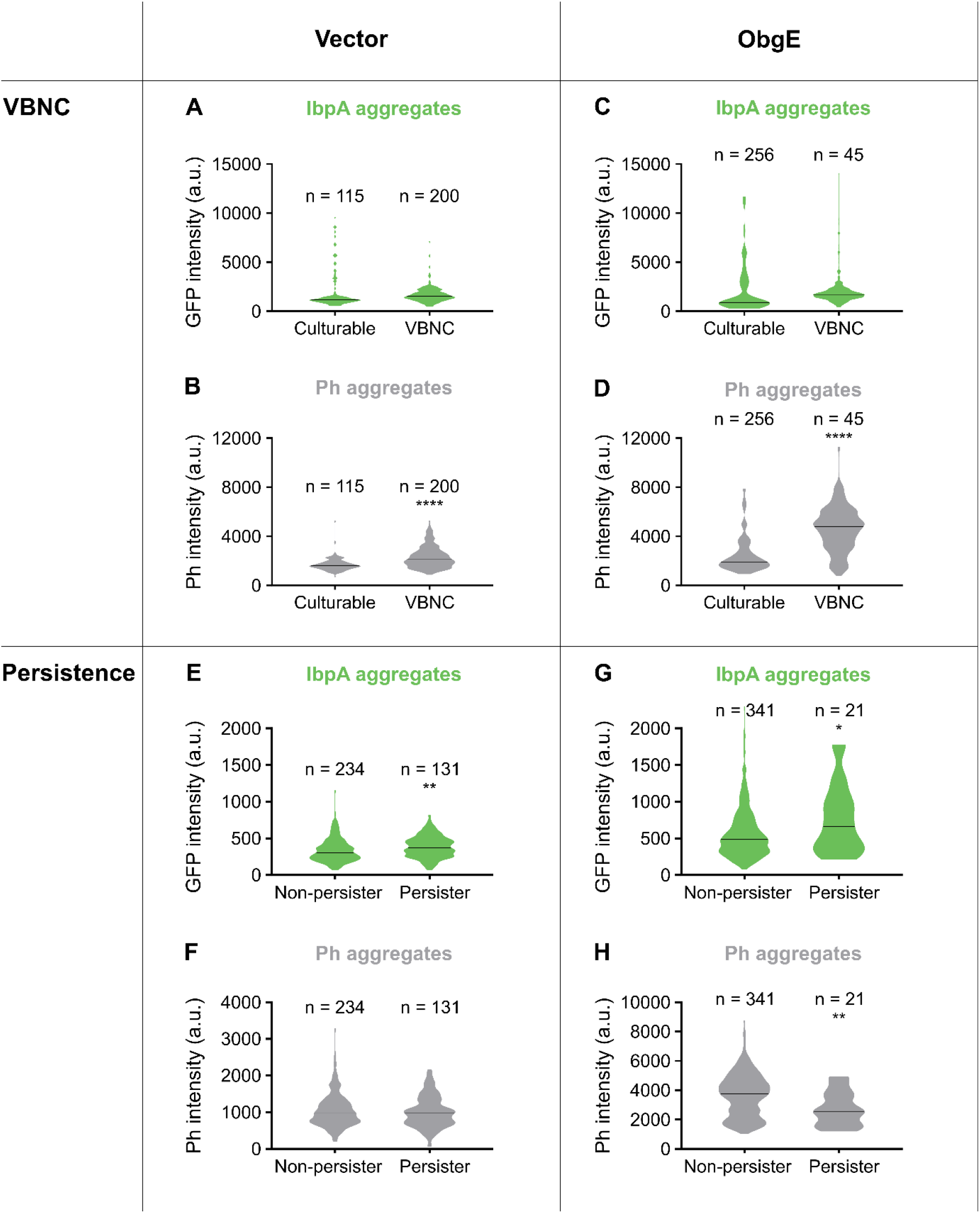
Bacterial dormancy, under the form of persistence and the VBNC state, is correlated with protein aggregation at the single-cell level. *E. coli ibpA-msfGFP* with pBAD33Gm (A and B) or pBAD33Gm-*obgE* (C and D) was grown for 24, 40 or 72 hours in the presence of the inducer arabinose. Cells were then immediately spotted on fresh medium to interrogate their resuscitation potential. Culturable cells were defined as cells that were able to resume cell division within 16 hours when provided with fresh nutrients. VBNC cells did not divide within this time frame. To assess persistence, *E. coli ibpA-msfGFP* with pBAD33Gm (E and F) or pBAD33Gm-*obgE* (G and H) was grown for 24 hours and treated with antibiotics before cells were spotted on fresh medium. Persister cells were defined as cells that were able to resume division within 16 hours after antibiotic treatment. The initial maximal IbpA-msfGFP fluorescence intensity of each cell (A, C, E and G) or their intensity in the phase contrast channel (B, D, F and H) is plotted as a measure for the extent of protein aggregation. Black lines indicate median values. n is the number of cells measured. * p < 0.05, ** p < 0.01, **** p < 0.0001.

**Figure S4:**
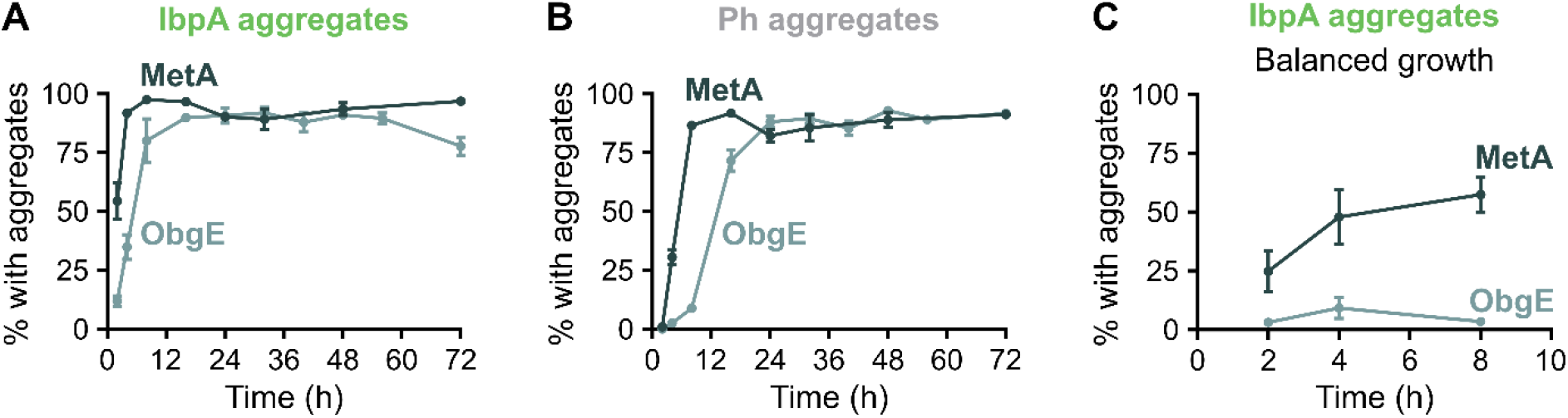
ObgE does not induce protein aggregation by acting as a nucleation factor. The percentage of cells of *E. coli* BW25113 *ibpA-msfGFP* overexpressing *obgE* (from pBAD33Gm) or *metA* (from pCA24N) that carry protein aggregates was determined by quantitative analysis of microscopy images. A-B) Cultures were allowed to enter stationary phase and aggregation was evaluated by the presence of fluorescent IbpA-msfGFP foci (A) and phase-bright structures (B) at several different time points. C) Cultures were maintained in balanced growth. At different time points, the percentage of cells that carry IbpA aggregates was determined. For every repeat and every time point at least 50 cells were analyzed. Data are represented as averages ± SEM, n ≥ 3.

**Figure S5:**
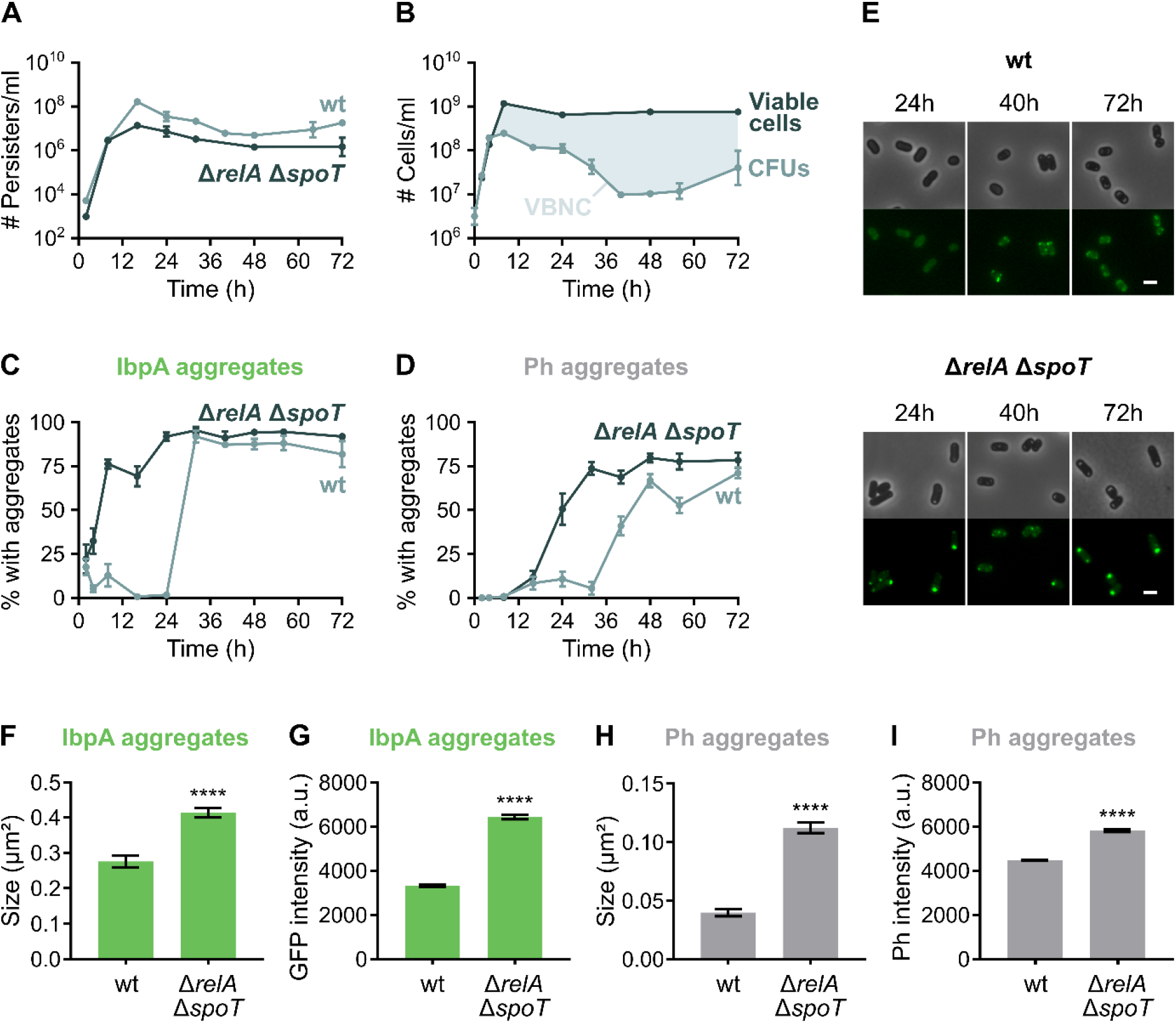
Protein aggregation, persistence and dormancy in *E. coli ΔrelA ΔspoT*. A) *obgE* overexpression leads to lower persister levels at almost all time points in the *ΔrelA ΔspoT* mutant in comparison to the wildtype. B) Quantification of CFUs and the total number of viable cells of *E. coli ΔrelA ΔspoT* upon *obgE* overexpression reveals that cells massively enter the VBNC state. Data are represented as averages ± SEM, n ≥ 3. C-D) The percentage of cells of a wild-type and *ΔrelA ΔspoT* culture that carry protein aggregates (in the absence of *obgE* overexpression) was determined by quantitative analysis of microscopy images. Aggregation was evaluated by the presence of fluorescent IbpA-msfGFP foci (C) and phase-bright structures (D). For every repeat and every time point at least 50 cells were analyzed. Data are represented as averages ± SEM, n ≥ 3. E-I) Protein aggregates that develop in *E. coli ΔrelA ΔspoT* (in the absence of *obgE* overexpression) are much larger and more intense than in the wildtype. This is shown by representative microscopy images (E), and quantitative measurements at time point 72h of the size (F) and intensity (G) of fluorescent IbpA-msfGFP foci and the size (H) and intensity (I) of phase-bright structures. Scale bar, 2 μm. Data are represented as averages ± SEM, n (number of cells analyzed) > 450 (**** p < 0.0001).

**Figure S6:**
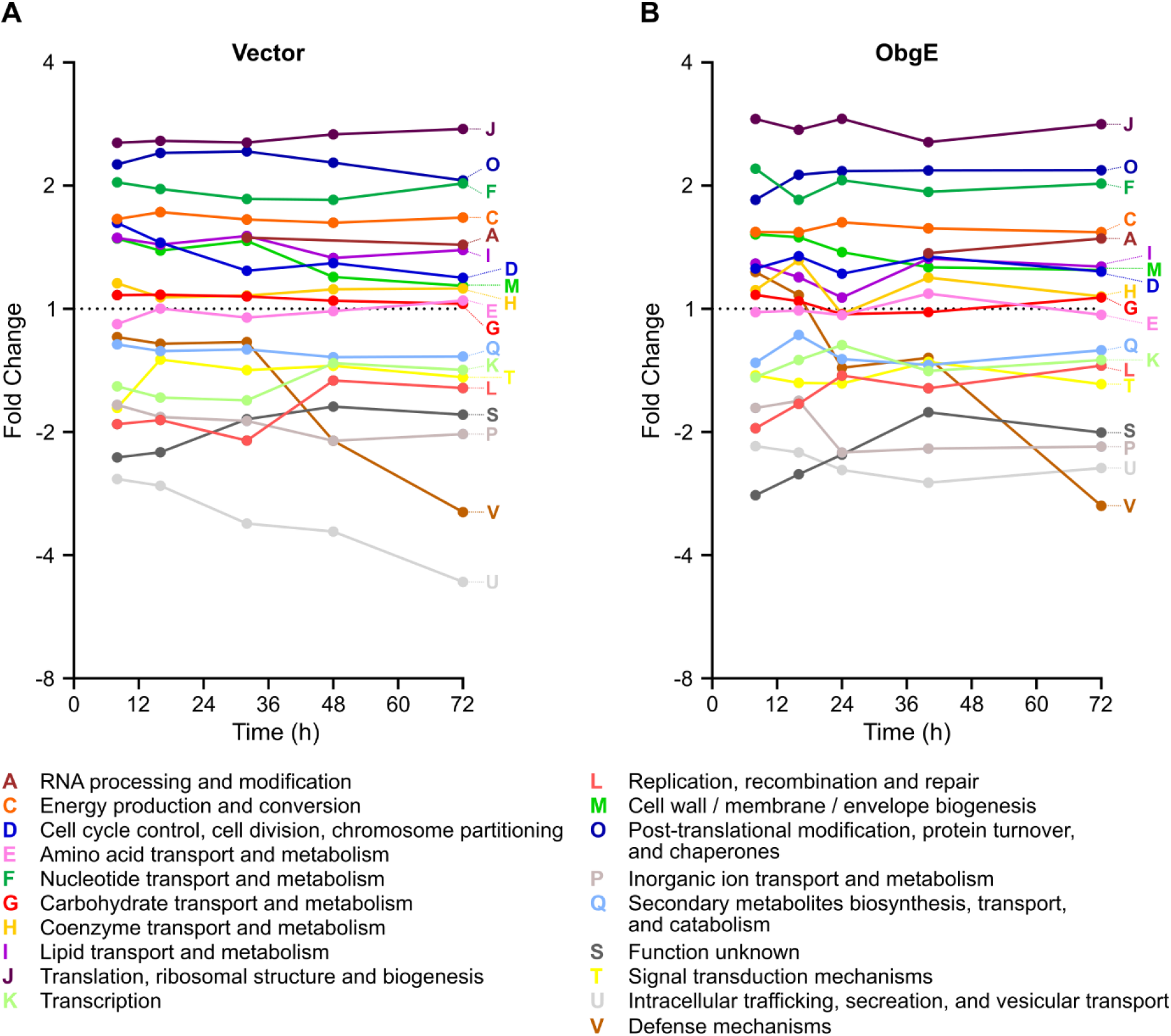
Biochemical analysis of isolated protein aggregates. A-B) Protein aggregates isolated from *E. coli* cultures carrying an empty expression vector (A) or overexpressing *obgE* (B) were subjected to MS analysis and COG functional categories were assigned to all identified proteins. The level of enrichment of every identified COG category is shown as a function of time. Significantly enriched categories are: C, F, J and O at all time points, M at the first time point for the vector and the second time point for ObgE. All other categories were never found to be significantly enriched (p < 0.05).

**Supplemental table 3:**
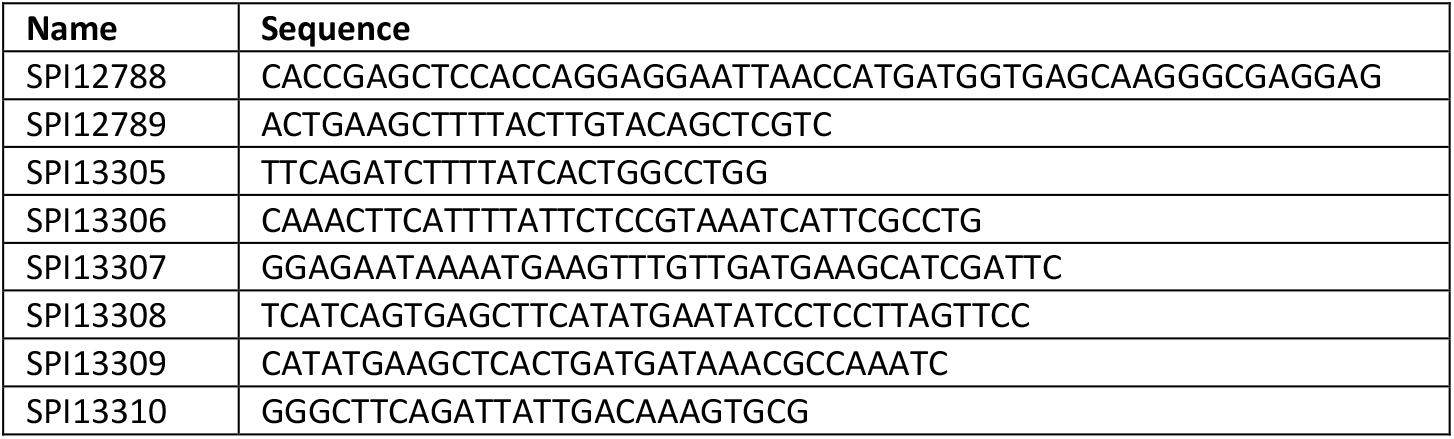
**Primer sequences.**

